# *Chlamydia gallinacea*: genetically armed as a pathogen however a phenotypical commensal?

**DOI:** 10.1101/2020.11.30.405704

**Authors:** Marloes Heijne, Martina Jelocnik, Alexander Umanets, Michael S.M. Brouwer, Annemieke Dinkla, Frank Harders, Lucien J.M. van Keulen, Hendrik Jan Roest, Famke Schaafsma, Francisca C. Velkers, Jeanet A. van der Goot, Yvonne Pannekoek, Ad P. Koets

## Abstract

*Chlamydia gallinacea* is an obligate intracellular bacterium that has recently been added to the family of *Chlamydiaceae*. *C. gallinacea* is genetically diverse, widespread in poultry and a suspected cause of pneumonia in slaughterhouse workers. In poultry, *C. gallinacea* infections appear asymptomatic, but studies about the pathogenic potential are limited. In this study two novel sequence types of *C. gallinacea* were isolated from apparently healthy chickens. Both isolates (NL_G47 and NL_F725) were closely related to each other and showed 99.1% DNA sequence identity to *C. gallinacea* Type strain 08-1274/3. To gain further insight in the pathogenic potential, infection experiments in embryonated chicken eggs and comparative genomics with *Chlamydia psittaci* were performed. *C. psittaci* is an ubiquitous zoonotic pathogen of birds and mammals, and infection in poultry can result in severe systemic illness. In experiments with embryonated chicken eggs *C. gallinacea* induced mortality was observed, potentially strain dependent but lower compared to *C. psittaci* induced mortality. Comparative analyses confirmed all currently available *C. gallinacea* genomes possess the hallmark genes coding for known and potential virulence factors as found in *C. psittaci* albeit to a reduced number of orthologues or paralogs. The presence of (potential) virulence factors and the observed mortality in embryonated eggs indicates *C. gallinacea* should rather be considered as a (conditional) pathogen than an innocuous commensal.

**Importance:** *Chlamydiaceae* are a family of bacteria comprising human and animal pathogens including the recently recognized *Chlamydia gallinacea. C. gallinacea* is widespread in poultry without causing clinical signs, which raises questions about its pathogenic potential. To assess this potential, two novel *C. gallinacea* strains were isolated, tested in infection experiments in embryonated chicken eggs and compared to *C. psittaci. C. psittaci* infection in poultry can result in severe systemic illness, depending on the conditions, and infections can be transmitted to humans. In the experiments *C. gallinacea* infection induced mortality of the embryo, but to a lower extent than infection with *C. psittaci*. Subsequent genome comparisons confirmed both *C. gallinacea* strains possess potential virulence genes typical for chlamydia, but fewer than *C. psittaci*. These results indicate *C. gallinacea* does have a pathogenic potential which warrants further research to elucidate its role as a poultry pathogen.

## Introduction

*Chlamydiaceae* are a family of obligate intracellular bacteria containing one genus and 14 species, and comprising human and animal pathogens. In birds, infections are caused by *Chlamydia psittaci* or more recently recognized species such as *C. gallinacea*. *C. psittaci* is zoonotic and has been reported worldwide in more than 465 bird species belonging to at least 30 orders. Most human infections have been linked to contact with birds or their environments (1). *C. gallinacea* is mainly detected in poultry with reports from almost all continents (2–4). *C. gallinacea* has incidentally been found in wild birds and cattle as a possible result of infection spill-over (5, 6). Possible zoonotic transmission of *C. gallinacea* has been considered but could neither be confirmed nor ruled out in slaughterhouse workers that were exposed to *C. gallinacea* infected poultry and developed pneumonia (7).

Infections with *C. psittaci* in birds are often asymptomatic, but can result in localized syndromes (e.g., conjunctivitis) or severe systemic illness. Chlamydial strain, avian host, host age and (environmental) stressors are important factors in the pathogenicity and occurrence of clinical signs (1). Studies investigating the pathogenesis of *C. gallinacea* are currently limited. As yet, clinical signs of disease in *C. gallinacea* infections have not been reported in observational field studies (2, 7, 8). Under experimental conditions it has been demonstrated that infection in broilers results in reduced weight gain (2). In a transmission study, *C. gallinacea* was mainly present in rectal and cloacal samples without clinical signs of disease and transmission occurred via the faecal-oral route (9).

Molecular studies using *ompA* genotyping or Multi Locus Sequence Typing (MLST) showed *C. gallinacea* is diverse, with at least 13 different *ompA* types and 15 different sequence types (ST) in 25 strains (2, 10). Fine detail comparative genomics revealed the *C. gallinacea* genome is conserved, syntenic and compact, but possess the hallmark of chlamydial specific virulence factors: inclusion membrane (Inc) proteins, polymorphic membrane proteins (Pmps), a Type III Secretion System (T3SS) and a plasticity zone with a cytotoxin (*tox*) gene (10, 11). Whether this genetic diversity and the presence of chlamydial virulence genes contributes to the pathogenicity of *C. gallinacea* remains a question, as clinical disease in infected chickens has not been reported in the limited number of field and experimental studies.

In this study, two novel *C. gallinacea* strains were isolated and compared to a virulent *C. psittaci* strain using an *in vivo* infection model in embryonated chicken eggs. In the eggs, *C. gallinacea* induced mortality was observed, but to a lower extent than *C. psittaci* induced mortality. In addition, molecular typing and comparative genomics with inter- and intra-species genomes was carried out. Both novel isolates represent different sequence types and possess the hallmark genes coding for known and potential virulence factors as found in *C. psittaci*, albeit to a reduced number of orthologs or alleles. The presence of these potential virulence factors and the observed mortality in *C. gallinacea* infected eggs indicates *C. gallinacea* should rather be considered as a (conditional) pathogen than an innocuous commensal.

## Results

### Isolation and pathology in embryonated eggs

Two novel strains (NL_G47 and NL_F725) of *C. gallinacea* were isolated and propagated in the yolk sac of embryonated chicken eggs. Strain NL_G47 was isolated from a caecal scraping sample collected in January 2018 from a 40-week old clinically healthy layer hen. Strain NL_F725 was isolated from a caecal suspension sample collected in August 2017 from a 34-week old layer hen. Both hens originated from different flocks, but were housed at the same location. Both flocks tested PCR positive for *C. gallinacea* in environmental boot sock samples about one month before the caecal samples were collected. The flock from which strain NL_F725 originated had to be culled to prevent spread of Infectious Laryngotracheitis (ILT). Three weeks before ILT infection was suspected the flock tested PCR positive for *C. gallinacea*. Background data of the flocks and a timeline are added to the supplementary (S1 and S2).

Primary isolation and propagation in Buffalo Green Monkey (BGM) cells failed, but after three passages in eggs the strains could be propagated in BGM cells. Replication of NL_G47 and NL_F725 was confirmed with positive immunofluorescence of the yolk sac membrane (see S3) and a positive *Chlamydiaceae* PCR targeting the 23S rRNA gene.

With the primary isolation of NL_G47, mortality was observed at day 10 after inoculation (incubation day 16) and at day 6 (incubation day 12) in the second passage. At primary isolation of NL_F725 no mortality of the embryos was observed, but eggs were harvested before day 10 after inoculation (day 8 after inoculation, incubation day 14) for logistical reasons. With the second passage of NL_F725, mortality of the embryos was observed at day 6 or day 7 after inoculation (incubation day 12 or 13). Based on egg candling, congestion of the blood vessels was observed prior to mortality of the embryos. At harvest the embryos were deep red (rubor), showed cyanotic toes and haemorrhaging of the skin (supplementary S3).

Histology and immunohistochemistry were performed to locate the bacteria in the different structures of the egg and to investigate any histological lesions. NL_G47 infected eggs were harvested at day 10 of incubation when anomalies of the vessels were observed with candling. Granular basophilic intracellular inclusions were seen in the epithelial cells of both the chorioallantoic membrane and the yolk sac membrane (Fig. 1A and C). These intracellular inclusions were strongly positive for chlamydial antigen labelling (Fig. 1B and D).

**Fig. 1.**
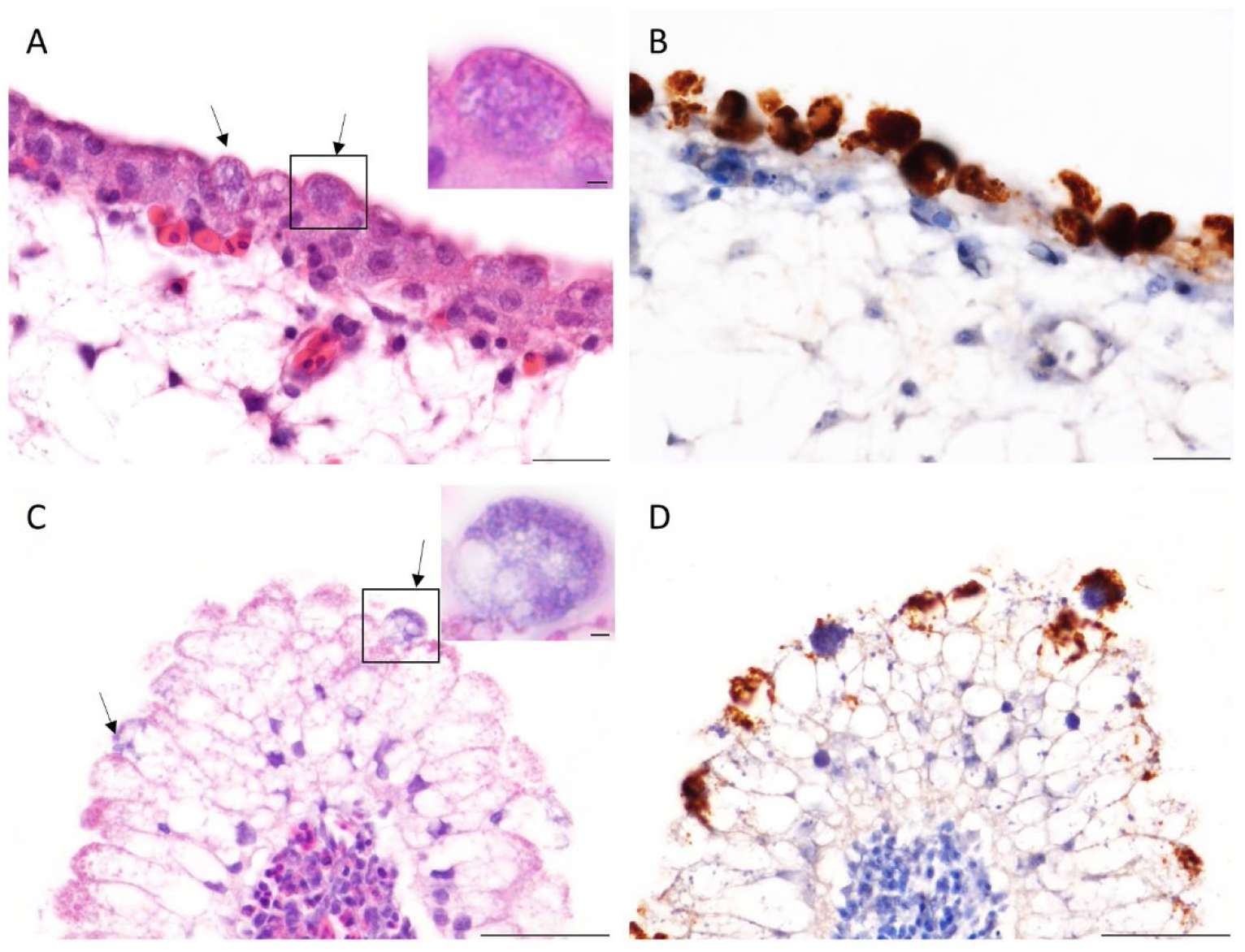
Chorioallantoic membrane and yolk sac membrane of 10 days embryonated eggs infected with NL_G47. Intracellular inclusions (arrows) in the epithelial cells of the chorioallantoic membrane (**A**) and yolk sac membrane (**C**). Inset: higher magnification showing the granular basophilic inclusions in the HE staining. Positive immunolabelling of the intracellular inclusions for chlamydial antigen in the chorioallantoic membrane (**B**) and yolk sac membrane (**D**). Scale bar is 20 micrometer (**A,B**), 50 micrometer (**C,D**) or 5 micrometer (insets).

### Assessment of virulence of *C. gallinacea* in embryonated eggs

Titration experiments in embryonated chicken eggs were performed to quantify the infectious dose and gain further insight in the pathogenic potential of the novel isolates compared to *C. psittaci*. Ten-fold serial dilutions of third passage yolk sac cultures of *C. gallinacea* NL_G47 and NL_F725, and *C. psittaci* NL_Borg, were used to calculate the egg infectious dose 50 (EID_50_) based on IFT positivity of the yolk sac membrane (with or without mortality of the eggs). As shown in Fig. 2A, the EID_50_ of *C. psittaci* strain NL_Borg was significantly higher than the EID_50_ (P<0.05, Wilcoxon-Mann-Whitney test) of *C. gallinacea* NL_G47. The EID_50_ of NL_F725 was in the same range as the EID_50_ of NL_G47, but could not be statistically assessed due the low number of observations.

**Fig. 2.**
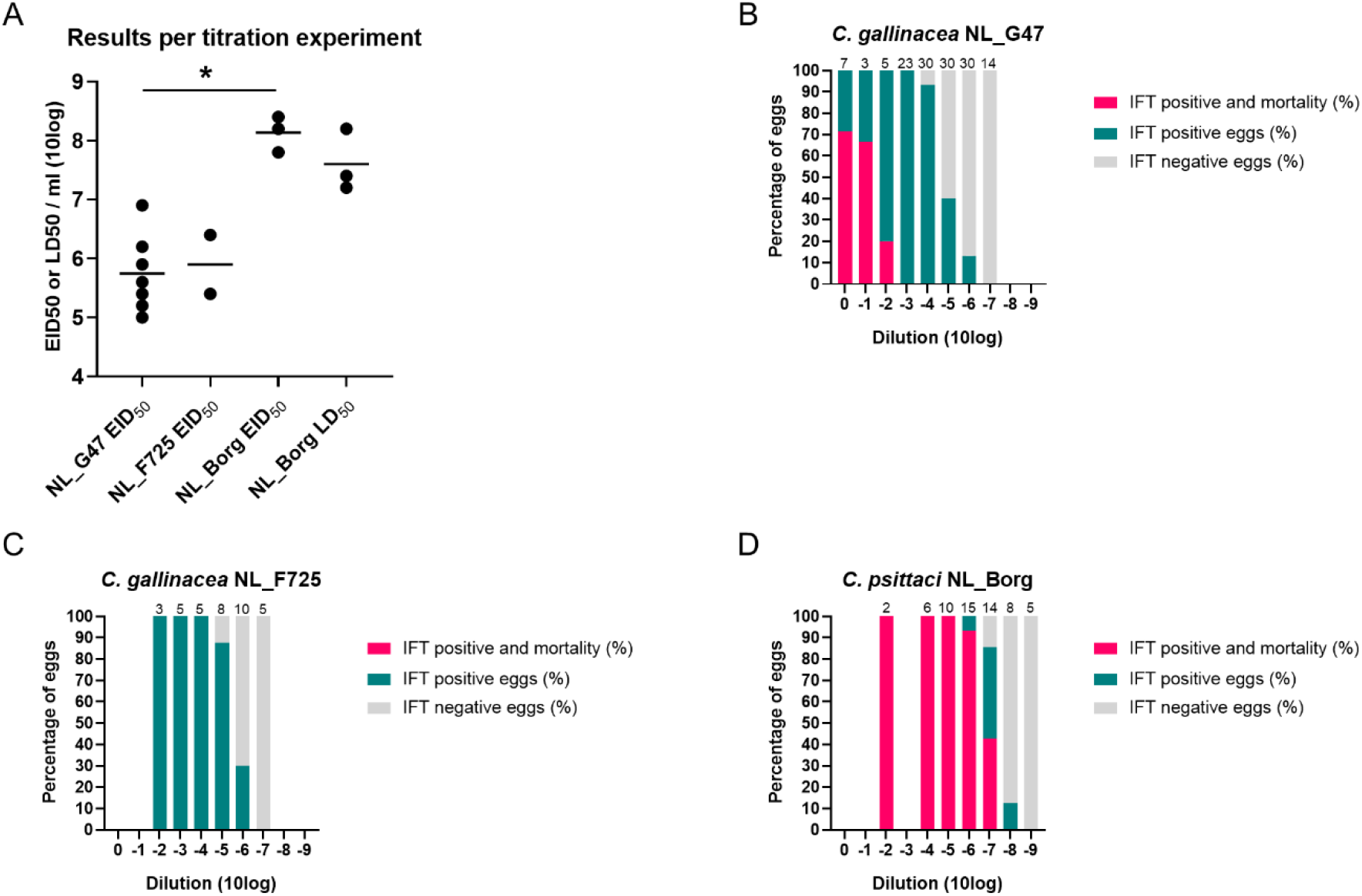
Assessment of virulence of *C. gallinacea* in embryonated eggs **A** shows the egg infectious dose 50 (EID50) of *C. gallinacea* NL_G47, NL_F725 and *C. psittaci* NL_Borg based on IFT of the yolk sac. The difference between EID50 of NL_G47 and NL_Borg was significantly different (*, P<0.05, Wilcoxon-Mann-Whitney test). For *C. psittaci* NL_Borg the lethal dose 50 (LD50) was also calculated. The median is indicated with a bar. **B, C** and **D** depict the cumulative results of the separate titration experiments per Chlamydia strain. Per dilution the percentage of eggs that was IFT positive with mortality, IFT positive without mortality and IFT negative are shown. The total number of eggs per dilution are presented at the top of every bar. These data are also included in the supplementary (S5).

For *C. psittaci* NL_Borg the lethal dose 50 (LD_50_) could also be calculated from the experiments. The LD_50_ of the experiments with *C. psittaci* NL-Borg showed overlap with the calculated EID_50_ (Fig. 2A). The LD_50_ from the experiments with *C. gallinacea* NL_G47 and NL_F725 could not be calculated because mortality was below the range of the dilution series that were used to calculate the infectious dose. To get further insight in differences in mortality and infectivity between *C. gallinacea* and *C. psittaci*, the egg data from all separate experiments were merged into one dataset (see supplementary S4).

The percentage of eggs that was IFT positive with mortality, IFT positive without mortality and IFT negative is shown per dilution and per *Chlamydia* strain (Fig. 2B-C-D). For strain NL_G47, mortality was observed until the 10^-2^ dilution and IFT positivity until the 10^-6^ dilution (Fig. 2B). For strain NL_F725 no mortality was observed in the dilutions that were tested (from 10^-2^ until 10^-7^), but IFT positivity was seen until the 10^-6^ dilution similar to strain NL_G47 (Fig. 2C). For strain NL_Borg, mortality was observed until dilution 10^-7^ and IFT positivity until 10^-8^ (Fig. 2D). These results indicate mortality in the *C. psittaci* infected eggs was relatively higher than in the *C. gallinacea* infected eggs and there might be a difference in mortality between *C. gallinacea* strains, although the number of observations was low.

### General characteristics of the genome sequences of Dutch *C. gallinacea* isolates

After isolation in eggs and one passage in BGM cells, DNA of both isolates was sequenced to identify their genetic background. The genomes of NL_G47 and NL_F725 have a total length of 1,066,007 and 1,064,097 bp, respectively, and include the chromosome and a plasmid (S5). Ribosomal MLST (rMLST) confirmed that both isolates belong to *C. gallinacea* (Fig. 3A), whilst the MLST showed that both isolates are genetically diverse and denoted with unique sequence types (ST280 and ST284). Phylogenetically, these clustered in distinct clades with NL_G47 forming a well-supported clade with the French isolate 08-1274/3, whilst NL_F725 clustered in a genetically diverse clade consisting of Chinese *C. gallinacea* strains (Fig. 3B).

**Fig. 3.**
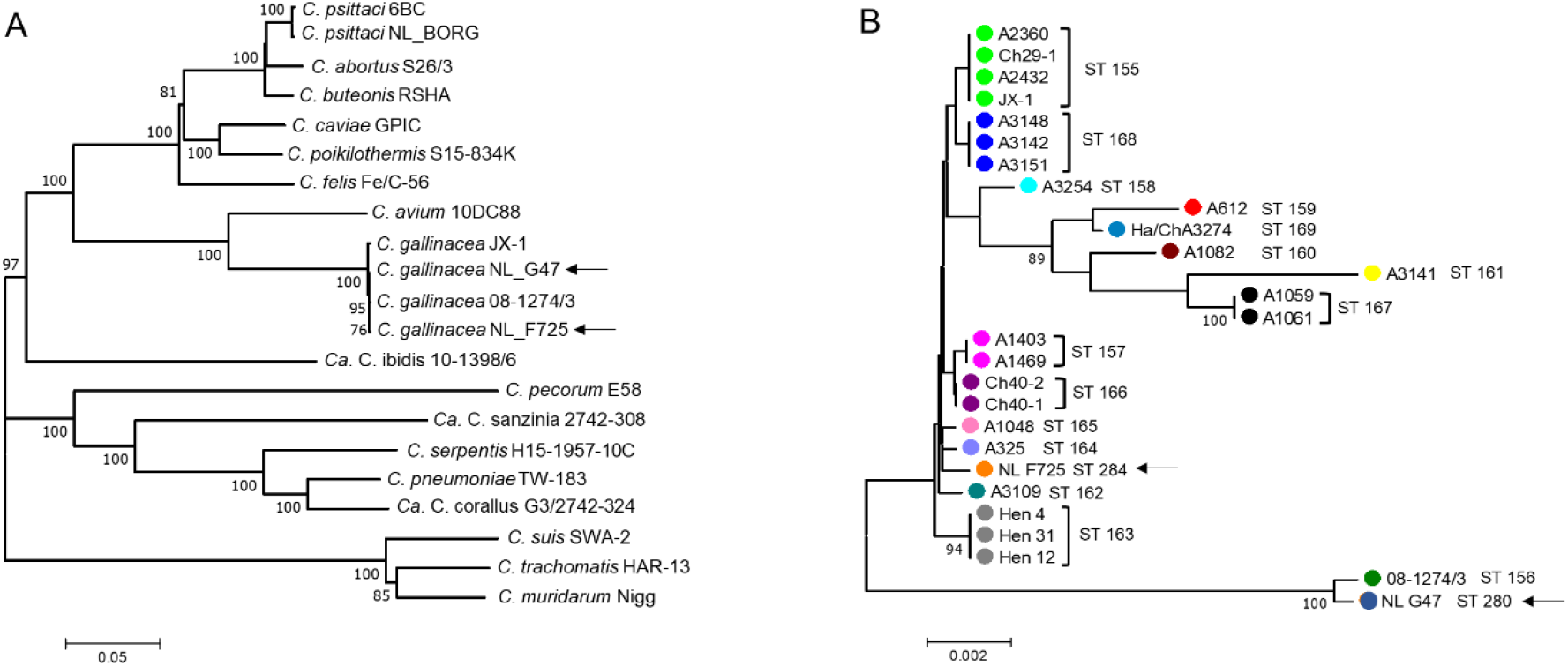
Phylogenetic analyses of concatenated sequences of *Chlamydia*. Concatenated sequences were aligned and analysed in MEGA7 (12). Phylogenetic trees were constructed by the Neighbour-Joining algorithm using the Maximum Composite Likelihood model. Bootstrap tests were for 1000 repetitions (13–15). Numbers on tree nodes indicate bootstrap values over 75% of the main branches. Horizontal lines are scale for genetic distances. **A** Neighbour-Joining tree of concatenated sequences of 52 ribosomal genes (rMLST)(16) of *Chlamydia* Type strains as well as three Candidatus (*Ca*. C. corallus, *Ca*. C. ibidis and *Ca*. C. sanzinia), *C. psittaci* strain NL_Borg and three additional *C. gallinacea* strains. All *C. gallinacea* strains (Dutch strains indicated by an arrow) clustered together in a well-supported and distinct clade with Chlamydia avium as the closest relative. **B** Neighbour-Joining tree of concatenated sequences of 7 housekeeping genes fragments (MLST)(17) of 27 *C. gallinacea* strains. Shared Sequence type (ST) in clades are indicated by color and ST number noted after brackets. Dutch *C. gallinacea* strains are indicated by an arrow.

### Comparative genome analysis of *C. gallinacea* and *C. psittaci*

To investigate genomic differences that might be related to the observed differences in the degree of pathology and mortality in eggs, the *C. gallinacea* and *C. psittaci* genomes were analysed and compared. *C. gallinacea* genomes NL_G47 and NL_F725 show approximately 100% synteny in genome organization and 99.1 % sequence similarity with *C. gallinacea* strains 08-1274/3 (type strain) and JX-1. All *C. gallinacea* genomes contain conserved hallmark chlamydial virulence genes coding for Incs, Pmps, T3SS and a PZ with a gene coding for the large cytotoxin (*toxB*). Most sequence variation was found in several distinct chromosomal regions and loci, namely in genes encoding the membrane proteins (e.g. *ompA* and *pmp*s), a conserved hypothetical protein, a phage tail protein, heme (*hemE*, and *hemN*) and glycogen (*glgP*) metabolism genes (S6). The PZ, a region of high genetic variability in chlamydial species, was conserved among the four *C. gallinacea* genomes with 99.3 – 99.8 % nucleotide identity (Fig. 5).

The genome sequence of our in-house reference strain *C. psittaci* NL_Borg was almost identical to reference strain *C. psittaci* NJ1 with only 65 Single Nucleotide Polymorphisms (SNPs). Due to the confirmed synteny in genome organization and high sequence similarity, type strain 08-1274/3 and NJ1 were used as representatives for *C. gallinacea* and *C. psittaci* species, in a translated coding sequences (CDSs) comparison. In the whole genome alignment, it was observed that the *C. psittaci* genome is 101.85 Kbp longer than the genome of *C. gallinacea* and contains more CDSs (Fig. 4A).

**Fig. 4.**
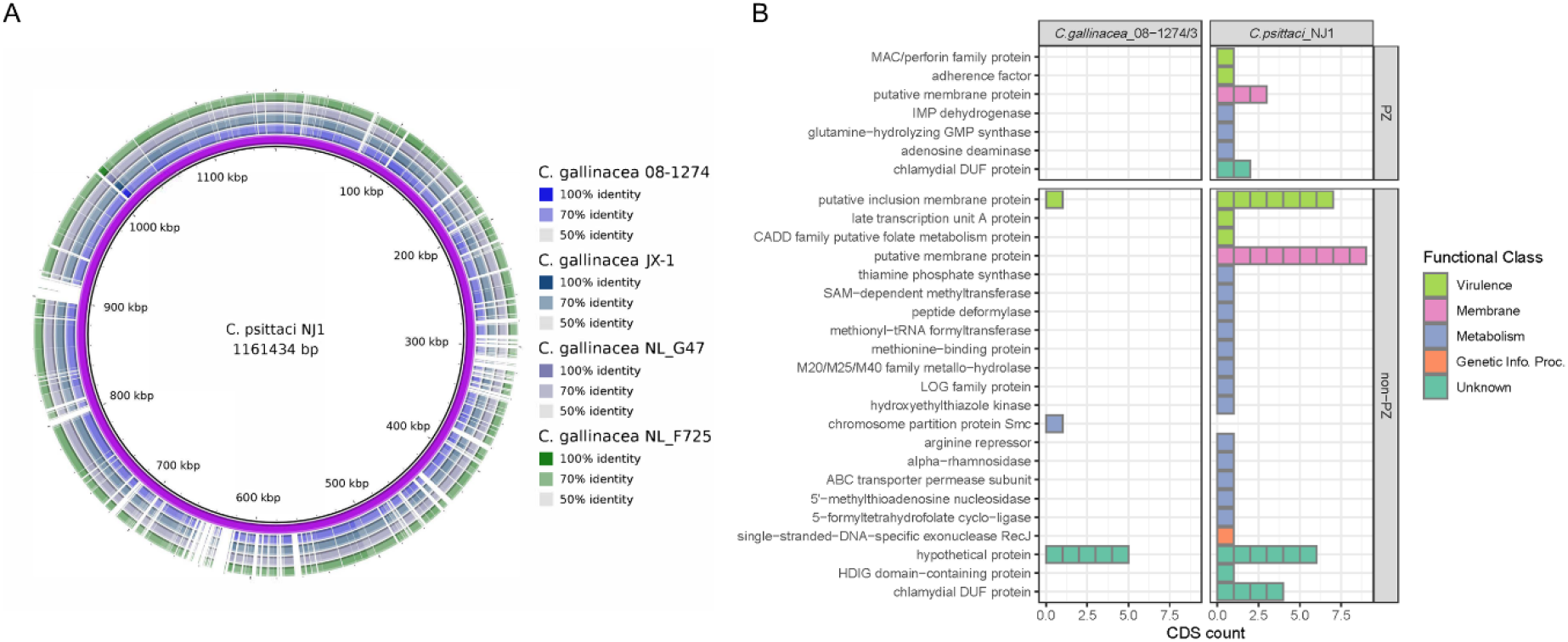
Genome comparison of *C. gallinacea* and *C. psittaci* **A** Whole genome BLAST comparisons between *C. psittaci* NJ1 and available C. gallinacea genomes created with BLAST Ring Image Generator (BRIG)(18). **B** CDS for which no homologue could be identified in *C. gallinacea* or *C. psittaci*. The proteins are categorized according to their function and location.

**Fig. 5.**
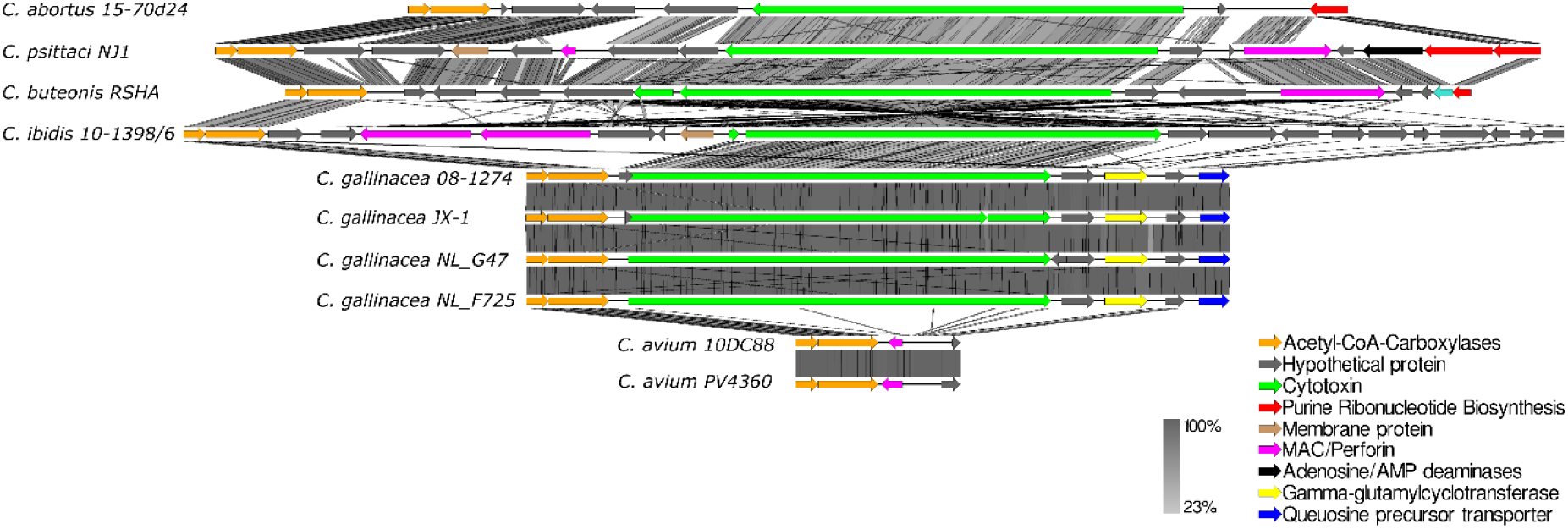
Graphical representation of the gene content of the PZ’s of representative Chlamydia species of avian origin including the Dutch *C. gallinacea* strains analysed in this study. Coloured arrows in the legend represent PZ genes according to function. Grey shading scale denotes % sequence similarity. The image was created with Easyfig (19).

With a local alignment approach, all translated CDSs of *C. gallinacea* 08-1274/3 (n=913) and *C. psittaci* NJ1 (n=986) and vice versa were compared to each other to identify regions with less or no homology (S7). The plasmids of *C. gallinacea* and *C. psittaci* were not included, because they are syntenic with both eight CDSs encoding the conserved chlamydial plasmid proteins.

As expected in closely related species, the majority of CDSs have orthologues in both species. In *C. gallinacea*, for only seven CDSs an orthologue could not be identified in *C. psittaci* (Fig. 4B, S8). Of those, one belonged to the family of putative Incs, a second had a metabolic function related to chromosome partition and the remaining five were hypothetical proteins with unknown function. In *C. psittaci*, 53 CDS could not be identified in *C. gallinacea*. Ten of these CDSs were located at the PZ coding for proteins such as the Membrane Attack Complex/Perforin domain-containing protein (MAC/PF), proteins involved in purine metabolism (*gua*AB-ADA operon) and a putative membrane protein. Although the length of the PZ of *C. gallinacea* is reduced compared to *C. psittaci*, it does contain an intact CDS for the cytotoxin (*toxB*), in contrast to the PZ of *C. avium* that lacks this gene (Fig. 5).

Outside the PZ, 18 of the unique CDS of *C. psittaci* were related to potential virulence factors (Fig. 4B). Most of these proteins belonged to the family of putative Inc proteins and membrane proteins. The remaining CDS were related to metabolism or to CDS coding for proteins of unknown function. Additional analysis of secretion signals of T3SS effector CDSs, important in *Chlamydia* virulence, revealed that a serine protease referred to as CPAF is not predicted to be excreted in *C. gallinacea* in contrast to *C. psittaci* (21). However, *C. psittaci* orthologues of the recently described T3SS that associate with the host’s inner nuclear membrane (SINC), and translocated actin-recruiting phosphoprotein (TARP) were identified and predicted to be secreted (S9).

Overall, the analysis revealed the novel *C. gallinacea* genomes NL_G47 and NL_F725 have 99.1 % sequence identity to the Type strain 08-1274/3 and include the hallmark chlamydial virulence genes. However, *C. psittaci* has a larger set of genes that are related to virulence and metabolism, including more *inc*s, *pmp*s, T3SS effectors and additional genes in the PZ.

## Discussions

In this study the pathogenicity and genetic background of two new chicken derived *C. gallinacea* strains (NL_G47 and NL_F725) were investigated combining classical methods with embryonated chicken eggs and a modern approach using whole-genome bioinformatic analyses. During isolation of NL_G47 and NL_F725 pathogenic changes, such as deep red colour (rubor), cyanotic toes and skin haemorrhage of the embryo, have been described for other *Chlamydia* species (22). Mortality in embryonated eggs after yolk sac inoculation with *C. gallinacea* has been reported by Guo et al. (2), but was not mentioned by Laroucau et al. (7).

The layer flocks from which the strains originated were apparently healthy, which is in line with observations from other field studies (2, 7, 8). It could not be evaluated if *C. gallinacea* infection led to impaired production as data on egg production were not collected in this teaching flock. The duration and frequency of shedding during *C. gallinacea* infection was only assessed to a limited extent due to the sampling strategy.

In the flock of strain NL_F725 the *C. gallinacea* infection preceded an infection with Infectious Laryngotracheitis (ILT) resulting in preventive culling to limit the spread of ILT. Whether a primary infection of *C. gallinacea* enhances infection with other pathogens or whether co-infection might exacerbate the disease outcome, is currently unknown. For *C. gallinacea*, only co-infections with *C. psittaci* have been reported in chickens without details about the clinical outcome (3, 23). For *C. psittaci* it has been suggested that co-infections with respiratory pathogens might lead to a more severe disease outcome (24, 25). The effect of co-infection could be a topic for future investigations.

In titration experiments in embryonated eggs, the pathogenicity of *C. gallinacea* was compared to a virulent *C. psittaci* poultry strain. The infectious dose and mortality in *C. gallinacea* infected eggs was lower compared to *C. psittaci* infected eggs. Furthermore, although the observations were limited, a small difference in pathogenicity between both *C. gallinacea* strains was observed. *C. gallinacea* NL_G47 infection resulted in mortality up to the 10^-2^ dilution (1 of 5 eggs), while no mortality was observed in the 10^-2^ dilution with strain NL_F725 (0 of 3 eggs). This is a first indication of a possible difference in pathogenicity between genetically different *C. gallinacae* strains, but needs to be confirmed due the low number of observations.

Furthermore, a higher mortality in *C. psittaci* infected eggs compared to *C. gallinacea* is in line with findings in available field and experimental studies. In these studies, *C. gallinacea* infection led to reduced weight gain in chickens and the absence of clinical symptoms, while exposure to a known high virulent *C. psittaci* strain can lead to severe systemic infections in chickens and turkeys (2, 8, 26, 27). In contrast, exposure to a less virulent *C. psittaci* strain resulted in mild respiratory symptoms indicating the importance of detailed strain knowledge and infection conditions (26).

The difference in infectious dose and mortality between *C. gallinacea* and *C. psittaci* in embryonated eggs might be a result of a shorter development cycle of *C. psittaci*. The development cycle of *C. gallinacea* takes about 60 to 72 hours while that of *C. psittaci* about 50 hours (1, 28). In the experiments all eggs were harvested at the same time point, which could mean *C. psittaci* was able to replicate to a higher number of bacteria. The difference in replication time could therefore contribute to the virulence of *C. psittaci*.

To get further insight in the genetic background of *C. gallinacea* in relation to pathogenicity, additional genomic comparisons were performed. Both *C. gallinacea* isolates were 99.1% identical to *C. gallinacea* Type strain 08-1274/3, with genetic diversity contained to several distinct chromosomal regions, and had a smaller set of potential virulence genes compared to *C. psittaci*. However, the question remains if a smaller set of virulence genes is a disadvantage for the particular isolate or species involved and determines the observed difference in pathogenicity. The closest genetical relative of *C. gallinacea*, *C. avium*, also has a reduced set of virulence genes compared to *C. psittaci*, and exhibits the smallest PZ region of all *Chlamydia*, but in cases involving pigeons and psittacines infection does lead to clinical signs and mortality (29, 30).

Moreover, *C. gallinacea* does contain all hallmark virulence factors such as Incs, Pmps T3SS and an intact cytotoxin in the PZ. In addition, *C. gallinacea* has genes encoding for the well-known T3SS effectors TARP and SINC that play a role in the pathogenesis of *Chlamydia* spp. In *C. psittaci*, TARP influences the active uptake in the host cell and SINC targets the nuclear envelope where it is hypothesized to interact with host proteins that control nuclear structure, signalling, chromatin organization, and gene silencing (31, 32). Future studies need to confirm if both effectors are indeed secreted in *C. gallinacea* and with which host proteins they interact.

Based on our current results in embryonated eggs and the genomic comparisons, it is too early to conclude if *C. gallinacea* is a phenotypical commensal. Although less pathogenic than the *C. psittaci* strains of avian origin, *C. gallinacea* does possess the hallmark *Chlamydia* virulence genes and infection does lead to mortality in embryonated chicken eggs after yolk sac inoculation. Furthermore, there might be small differences in virulence between *C. gallinacea* strains. Additional pathogenesis studies in chickens, including predisposing conditions such as co-infections, are therefore needed to further elucidate the pathogenic potential of *C. gallinacea* and possible strain differences. These future studies will help to assess the importance of this pathogen for poultry industry.

## Methods

### Ethical statement

The cloacal and caecal sampling of the chickens was approved by the Dutch Central Authority for Scientific Procedures on Animals and the Animal Experiments Committee (permit number AVD108002016642) of Utrecht University (the Netherlands) and all procedures were conducted in accordance with national regulations on animal experimentation. No ethical approval is required for work with embryonated chicken eggs until day 18 according to Dutch Law.

### Biosafety

All culture work with *C. gallinacea* was performed under biosafety level 2 and all culture work with *C. psittaci* under biosafety level 3.

### Sample collection and inoculum preparation

Layer flocks at the Faculty of Veterinary Medicine in Utrecht, the Netherlands were monitored for the presence of *C. gallinacea* with boot sock sampling. The flocks were obtained from commercial laying hen rearing farms at 18-weeks of age and had an average size of 50 hens that were distributed evenly over two pens. Background data on the flock are supplied in S1 and S2. From each pen, environmental boot sock samples were collected monthly. When the boot socks turned PCR positive for *C. gallinacea*, individual cloacal swabs and caeca were collected. Swabs were stored in one millilitre Sucrose Phosphate Glutamate (SPG) and caeca in ten percent weight per volume (w/v) according to standard protocols (33, 34). SPG contains sucrose (75 g/litre), KH_2_PO_4_ (0.52 g/litre), K_2_HPO_4_ (1.25 g/litre) and L-glutamic acid (0.92 g/litre). Before use, fetal bovine serum (0.1 ml/ml), amphotericin B (4 μg/ml), gentamicin (40 μg/ml and vancomycin (25 μg/ml) were added. Samples were stored at −80 °C.

To prepare the inoculum for the eggs, swabs were thawed at room temperature for approximately one hour. Swabs were centrifuged for ten minutes at 500 x *g* and 200 microliter of the supernatant was used for inoculation. Caeca were prepared following two methods. For the isolation of NL_G47 the caecum was cut lengthways in parts of approximately 2 cm. Subsequently the parts were washed in SPG and the epithelium was removed by scraping with a scalpel. The scrapings of epithelium were washed in 2 ml of SPG and the suspension was filtered over a 0.8 μm filter (Acrodisc^®^ Syringe Filter, Pall Life Sciences). After 1 hour of incubation at room temperature the suspension was used for inoculation.

For the isolation of NL_F725, caeca were homogenized in a ten percent w/v suspension in an ULTRA-TURRAX tube (BMT-20-S, IKA) on an ULTRA-TURRAX^®^ Tube Drive (IKA) at 6000 RPM for 90 seconds and switching direction every 30 seconds. The suspension was centrifuged at 500 x *g* for 15 minutes and the supernatant was used for culturing.

### Isolation in eggs

#### Inoculation

Specific pathogen free (SPF) embryonated chicken eggs were delivered after five days of incubation, candled to check viability and incubated overnight at 37.5 – 38 °C and 65 % relative humidity in small egg incubators (Octagon 20 Advance, Brinsea). Inoculation was performed at day six of incubation (one day after delivery).

Before inoculation, the eggs were candled, and the air chamber was marked with a pencil. The eggs were cleaned with a wipe drenched in 70 percent ethanol. In the middle of the area of the marked air chamber, a hole was drilled with a 0.8 mm engraving bit (26150105JA, Dremel). Subsequently, the eggs were moved to a flow cabinet and sprayed with 70 percent ethanol. Per egg, 200 μl was inoculated in the yolk sac with a one millilitre syringe and a 22G x 40mm needle. The full needle was inserted perpendicularly into the drilled hole.

Per clinical sample, four eggs were inoculated. As a negative control, 2 eggs were inoculated with Dulbecco’s Phosphate Buffered Saline (DPBS, Gibco, Life Technologies Limited) and, as a positive control, 2 eggs were inoculated with *C. gallinacea* strain 08DC65. Strain 08DC65 was obtained from the Friedrich Loeffler Institute in Jena, Germany.

After inoculation eggs were wiped with 70 % ethanol and the hole was closed with a droplet of nail polish. The eggs were placed in the egg incubators and incubated until day 16 or until mortality. At day 16, eggs were chilled overnight at 4 °C for further processing.

#### Candling

Mortality was monitored by daily candling. With candling, the appearance of vessels and movement of the embryo was monitored (35). The result of candling was graded:

- no abnormalities observed: vessels are visible, movement of the embryo
- abnormalities observed: congestion or bleeding from vessels, decreased movement of the embryo
- mortality: no or less vessels visible and no movement of the embryo

When abnormalities were observed an extra check was performed on the same day. After mortality or an increase in the severity of the abnormalities, eggs were chilled overnight at 4 °C until harvesting.

#### Harvesting

Mortality within three days after inoculation (day nine of incubation) was considered as acute mortality inconsistent with a *Chlamydia* infection (22). These eggs were disinfected with 70 percent ethanol, opened at the air sac side and checked for any visual deformations. Furthermore, a sheep blood agar plate was inoculated with a loopful from the yolk sac and incubated overnight at 37 °C to check for bacterial contamination.

Eggs were harvested for the isolation of *C. gallinacea* when mortality occurred from day nine of incubation or when no mortality was observed at day 16 of incubation. At harvesting the part of the egg shell covering the air sac was removed, and subsequently the egg shell membrane and the allantois membrane were opened with disposable tweezers. The allantoic fluid was removed with a pipette, the egg was then emptied in a Petri dish to harvest the yolk sac membrane. The yolk sac membrane was weighted and transferred to an ULTRA-TURRAX tube (BMT-20-S, IKA). Depending on the volume of the yolk sac and the size of the tube, SPG buffer was added and the yolk sac membrane was homogenized on an ULTRA-TURRAX^®^ Tube Drive (IKA) during 90 seconds (switching between forward and reverse every 30 seconds) at 6000 RPM. The suspension was transferred to 50 ml Falcon tubes and SPG buffer was added until a 20 % w/v suspension.

The yolk sac membranes from eggs inoculated with the same sample and harvested at the same day were pooled. A 10 μl droplet of the yolk sac suspension was spotted in duplo on glass slides and air dried. The glass slides were tested with the IMAGEN^tm^ Chlamydia test kit according to manufacturer’s instructions (Thermo Scientific). Two hundred μl of the suspension was used for PCR testing.

### Titration experiments in embryonated eggs

The isolated strains and *C. psittaci* strain NL_Borg were tested in titration experiments. The experiments were repeated 7 times for NL_G47, 2 times for NL_F725 and 3 times for NL_Borg. Strain NL_Borg was selected because it is genetically closely related to strain FalTex and NJ1, which are both isolated from outbreaks in poultry (turkeys) (36).

To standardise the inocula before the experiments all three strains were passaged three times in embryonated eggs under similar conditions. The third passage yolk sac membrane suspensions were used to prepare tenfold serial dilutions in DPBS (Gibco, Life Technologies Limited) for inoculation of the yolk sac of 6-day incubated chicken eggs. The eggs were incubated at 37 °C and 65% relative humidity in egg incubators (Octagon 20 Advance, Brinsea). After mortality or 6 days after inoculation the eggs were chilled overnight at 4 °C and harvested as described earlier.

In a first experiment the range for the dilution series was defined by inoculating a limited number of eggs per dilution. In a subsequent experiment the range was limited to four dilution steps. Per dilution step four or five eggs were inoculated with 200 μl suspension. Two eggs were inoculated with sterile DPBS (Gibco, Life Technologies Limited) as a negative control and, as a positive control, 2 eggs were inoculated with a lower dilution of the *Chlamydia* strain that was used in the experiment.

After each titration experiment the 50% egg infectious dose (EID50) and, when possible, the 50% egg lethal dose (LD50) per ml inoculum was calculated according the Spearman-Karber method (37, 38). The difference in EID50 between strains was assessed using the Wilcoxon-Mann-Whitney test.

### Isolation in cell culture

Isolation and propagation in cell culture was performed as described earlier (29). Briefly, Buffalo Green Monkey (BGM) cells were seeded with Dulbecco ‘s Modified Eagle Medium (DMEM, Gibco, Life Technologies Limited) and 10% serum in 24-well plates (Greiner Bio-One GmbH, Germany). The plates were incubated at 37 °C with 5% CO_2_ in a humidified incubator until 80% confluency of the monolayer. After inoculation, the plates were centrifuged at 2450 × g and 37 °C for 60 min and subsequently incubated for two hours. The medium was then replaced with UltraMDCK serum-free medium (Lonza). At day 1 and day 4, 200 μl of the supernatant was collected for PCR to monitor replication. Plates were harvested at day 4 for DNA isolation, further passaging or storage at −80 °C.

### Histology and immunohistochemistry

From infected and non-infected eggs the chorioallantoic membrane, yolk sac and embryo were harvested for histology and immunohistochemistry. After fixation in 10% neutral buffered formalin, tissues were routinely processed into paraffin blocks. Four micrometer sections were cut and collected on coated glass slides. Sections were stained with haematoxylin-eosin (HE) or immuno-stained with a polyclonal anti-Chlamydia antibody (LS-C85741) and a monoclonal anti-Chlamydia antibody (MBS830551).

For the polyclonal antibody the antigen was retrieved by proteolysis-induced epitope retrieval (0.1% Trypsin in TBS for 30 min at 37 °C). For the monoclonal antibody heat-induced epitope retrieval was used (citrate buffer, pH 6.0, 21°C for 5 min). The primary antibody (dilution 1:100) was incubated for 60 minutes. HRP EnVision anti-Mouse or HRP Envision anti-Rabbit (Dakopatts) were used as a secondary antibody for 30 min, depending on the nature of the first antibody. Subsequently, sections were incubated for 5 minutes in DAB+ substrate (Dakopatts) and then counterstained with Mayer’s haematoxylin.

### Molecular techniques

#### PCR

Two hundred μl of the washing suspension, yolk sac suspension or cell culture supernatant was used for DNA isolation. DNA isolation was performed with a MagNA Pure LC total Nucleic Acid Isolation kit in the MagNA Pure^®^ system (Roche Diagnostics, Almere, the Netherlands). Samples were tested with a *Chlamydiaceae* PCR targeting the 23S rRNA and *C. gallinacea* PCR targeting the *enoA* gene or *C. psittaci* PCR targeting the *ompA* gene as described earlier (8, 39).

#### Genome Sequencing, Assembly, Annotation and Mapping

Twenty-four-well cell culture plates were freeze-thawed twice and the cells were subsequently harvested for DNA isolation as described earlier (29). DNA was isolated according to the DNeasy Blood and Tissue kit (Qiagen GmbH, Germany).

The DNA samples were prepared for Illumina sequencing using the SMARTer^®^ ThruPLEX^®^ DNA-Seq kit (Takara Bio, USA) according to manufacturer protocol. Quality control of the library preparation was performed on a Tapestation 2200 (Agilent Technologies, Germany) and the DNA concentration was determined on a Clariostar (BMG Labtech, the Netherlands) with use of the Quant-IT PicoGreen^®^ dsDNA kit (Invitrogen Ltd, UK). Sequencing was performed on an Illumina MiSeq platform. The complete genome and plasmid sequences were assembled using SPAdes 3.9 (40). Contigs containing sequences of BGM cells were removed prior to subsequent analysis. Genome size was determined to be 114660 bp for NL_G47 (coverage depth 366), 1064097 bp for NL_F725 (coverage depth 181), 1167736 bp for NL_Borg (coverage depth 633). All strains had a plasmid size of 7500 bp.

Genomic DNA of isolates NL_G47 and NL_F725 was also used for long read sequencing on the Oxford Nanopore Technologies platform. Multiplex libraries were prepared using the Ligation sequencing Kit SQK-LSK109 according to the manufacturer’s protocol. Sequencing was performed on the MinION Mk-1C using a FLO-MIN106D flowcell. Nanopore sequences were cleared from adapters with Porechop v0.2.3_seqan2.1.1 and reads of low quality and shorter than 1000 bp were removed with NanoFilt v2.7.1 (41, 42).

To generate a closed genome for *C. gallinacea* isolate NL_F725 we used a hybrid assembly approach where contigs constructed from Illumina reads and Nanopore reads are combined. The hybrid assembly was generated using the Unicycler v0.4.8 in *conservative* mode (43).

Assembled contigs were annotated using the RGAP pipeline using a corresponding reference genome (44).

#### Accession numbers

Read data generated for this study, and the annotated assemblies for the strains, can be found in ….needs to be added…

The sequences are deposited into xxx and the public available Bacterial Isolate Genome Sequence Database (BIGSdb) ((http://pubmlst.org/chlamydiales) (*C. gallinacea* isolates NL_G47 (id: 4548) and NL_725 (id: 4560) and *C. psittaci* NL_Borg (id: 4561).

#### Molecular typing

Phylogenetic trees were generated by exporting gene sequences from the *Chlamydiales* database (http://pubmlst.org/chlamydiales) as an XMFA file containing each locus as an aligned block. The XMFA file was converted to an aligned concatenated sequence for neighbor-joining tree analysis using MEGA7(12).

For rMLST complete sequences (~22.000 bp) of 52 genes encoding ribosomal proteins (*rps*) were analysed (16). The *rps* gene *rpmD*, encoding the 50S ribosomal protein L30 is absent in genomes of *Chlamydia* isolates analysed so far.

For MLST, sequences of fragments (400 – 500 base pairs) from seven housekeeping genes (*enoA, fumC, gatA, gidA, hemN, hlfX, oppA*) were analysed (17). Isolates used for rMLST and MLST including provenance and allelic profile data are listed in the supplementary (S12 and S13).

##### Genome comparisons

Average nucleotide identity (ANI) determination for the newly sequenced *C. gallinacea* genomes was performed at enve-omics.ce.gatech.edu/ani/, whilst the genome completeness and quality of the assemblies was estimated using Quast (45–47). SNPs in contigs assembled from Illumina reads were identified using Snippy v4.6.0 (48).

*C. gallinacea* pairwise genome comparisons were performed using the Geneious Prime 2020.2 platform (https://www.geneious.com). The genomic regions of interest and/or polymorphic loci were extracted from the analysed genomes and aligned with MAFFT and/or Clustal Omega (as implemented in Geneious Prime) for further nucleotide and/or translated protein sequence analyses performed using DNASp 6.0 (49). The total number of polymorphisms (and gaps), % nucleotide and amino acid sequence identity, number of haplotypes and haplotype diversity (Hd), and ratios of the rates of non-synonymous to synonymous nucleotide substitutions per site (dn/ds) averaged over the entire alignment were calculated.

As the Type 3 Secretion System (T3SS) play a key role in the interaction of chlamydia and hosts, EffectiveDB (http://effectivedb.org) was used to predict the T3S secreted proteins of *C. gallinacea*. For prediction the standard Effective T3 classification module 2.0.1 was used with a cut-off score of 0.9999 (20). Similarly, to predict transmembrane *C. gallinacea* proteins, and identify inclusion membrane proteins characterised by bilobed hydrophobic domains, TMHMM 2.0 server (https://services.healthtech.dtu.dk/service.php?TMHMM-2.0) was used (50).

The visualisation of BLAST comparisons of our newly sequenced draft *C. gallinacea* genomes to published *C. gallinacea* genomes 08-1274/3 and JX-1, and/or *C. psittaci* NJ1 was performed with BLAST Ring Image Generator (BRIG) (18). Visualisation of the BLAST comparison, sequence identity and genomic structure of the plasticity zone for C*. gallinacea* and those from other related species was performed using EasyFig (19).

#### Comparative analyses of *C. psittaci* and *C. gallinacea* CDSs (translated proteins)

For the identification of orthologous genes in *C. gallinacea* and *C. psittaci*, an all-vs.-all comparison of the translated coding sequences (CDSs) was performed using global sequence alignment of each CDS. Translated CDSs were aligned using DIAMOND v0.9.14 and the best hit for each query was selected (51). Only hits with an expect (E) value less than 10^-3^ were included. CDS with no hits or hits with an E-value above the threshold were further investigated and the annotation artefacts were removed. The remaining CDS were assigned unique. Results of the alignment were structured and visualized using the *tidyverse* package and R v3.6.1 (52, 53).

## Acknowledgements

The authors acknowledge Herma Buys, Irene Oud and Marianne Vahl of the WBVR diagnostic lab and for their assistance with PCR tests; Marielle van den Esker for proof reading; Lars Ravesloot for optimizing pictures in the supplementary; Arie Kant and Quillan Dijkstra for technical assistance with Nanopore sequencing; the animal caretakers, Carmen Minnee, Freek Weites and Marc Kranenburg of the department Population Health Sciences, division Farm Animal Health of the faculty of Veterinary Medicine in Utrecht for their assistance in sampling the chickens. The authors would also like to thank Dr Christiane Schnee from the Friedrich Loeffler Institute in Jena, Germany for providing strain *C. gallinacea* 08DC65.

